# Transcriptomic dysregulations associated with SARS-CoV-2 infection in human nasopharyngeal and peripheral blood mononuclear cells

**DOI:** 10.1101/2020.09.09.289850

**Authors:** Caroline Vilas Boas de Melo, Maruf Ahmed Bhuiyan, Winfred Nyoroka Gatua, Stephen Kanyerezi, Leonard Uzairue, Priscilla Abechi, Karan Kumar, Jabale Rahmat, Abdulazeez Giwa, Gracious Mwandira, Abisogun Mujib Olamilekan, Tiffany Ezinne George, Oluwapelumi John Adejinmi, Monsurat Ademidun Ibironke, Olayemi David Rotimi, Dina Aly Mahmoud Aly Abo-Elenein, Ridwanullah Abiodun Abubakar, Mahmood Usman, Ifeoluwa Adewunmi, Oyewumi Akinpelu, Olajide Emmanuel, Khatendra Reang, Akadiri Olalekan, Sarah H. Carl

**Affiliations:** Laboratório de Patologia Estrutural e Molecular, Instituto Gonçalo Moniz, Fundação Oswaldo Cruz, Rua Waldemar Falcão, 121, Candeal – Salvador, Brazil. CEP: 40296-710; Department of Virology, Bangabandhu Sheikh Mujib Medical University, Shahbag, Dhaka, 1000, Bangladesh.; A. Department of Biochemistry and Biotechnology, Pwani University, P.O.BOX 195- 80108 KILIFI, Kenya; B. Molecular Biology and Bioinformatics Unit, International Centre of Insect Physiology and Ecology, P.O.BOX 30772-00100 NAIROBI, Kenya.; Department of Immunology and Molecular biology, College of Health Sciences, Makerere University. Kampala, Uganda, 7072.; Department of Microbiology, College of Bioscience, Federal University of Agriculture, Abeokuta, Ogun State, Nigeria, 3100001.; NGN, Biological sciences, African Center For Excellence In Genomics Of Infectious Diseases (ACEGID), Redeemer’s University Ede Osun state.; Centre for Energy, Indian Institute of Technology Guwahati, Guwahati, Assam- 781039, India.; School of Life Sciences Independent University, Bangladesh (IUB) Plot 16 Aftab Uddin Ahmed Rd, Dhaka 1229.; South African National Bioinformatics Institute, University of the Western Cape, South Africa.; University of Malawi, College of Medicine, Biomedical Sciences department, Private bag 360, Chichiri Blantyre 3.; Department of Cell Biology and Genetics (Genetics), University of Lagos, Lagos Nigeria.; Biotechnology, Federal University of Technology, Owerri (FUTO), Owerri Imo State Nigeria, 4130231.; College of Medicine, University of Ibadan.; Department of Botany, Lagos State University, KM 15, Badagry Expressway, Ojo, PMB 0001, LASU Post Office, Ojo, Lagos, Nigeria.; Biochemistry, Ekiti State University, Ekiti State University, PMB 5363, Iworoko road, Ado Ekiti, Ekiti State. Nigeria. Pincode: 362001.; Zoology Department, Cairo Science University.; Department of Physiology, University of Ilorin, Nigeria PMB 1515.; Anatomy Department, Ahmadu Bello University, Zaria – Nigeria.; Physiology, University of Ibadan, Nigeria.; University of Strathclyde, Glasgow, United Kingdom, G1 1XQ.; Nematology Research Unit, Department of Biology, Ghent University, K.L. Ledeganckstraat 35, 9000 Gent, Belgium.; Hislop College, Biochemistry Department, Chitnavis Centre, Temple Rd, Dobhi Nagar, Civil Lines, Nagpur, Maharashtra 440001.; Biotechnology, Federal Institute of Industrial Research Oshodi Lagos, Nigeria.; Scailyte AG, Industriestrasse 12, 6210 Sursee, Switzerland.

**Keywords:** SARS-CoV-2, COVID-19, Transcriptomic analysis, Nasopharyngeal swab, PBMC

## Abstract

**Introduction:** Over 24 million people have been infected globally with the novel coronavirus, SARS-CoV-2, with more than 820,000 succumbing to the resulting COVID-19 disease as of the end of August 2020. The molecular mechanisms underlying the pathogenesis of the disease are not completely elucidated. Thus, we aim to understand host response to SARS-CoV-2 infection by comparing samples collected from two distinct compartments (infection site and blood), obtained from COVID-19 subjects and healthy controls.

**Methods:** We used two publicly available gene expression datasets generated via RNA sequencing in two different samples; nasopharyngeal swabs and peripheral blood mononuclear cells (PBMCs). We performed a differential gene expression analysis between COVID-19 subjects and healthy controls in the two datasets and then functionally profiled their differentially expressed genes (DEGs). The genes involved in innate immunity were also determined.

**Results:** We found a clear difference in the host response to SARS-CoV-2 infection between the two sample groups. In COVID-19 subjects, the nasopharyngeal sample group indicated upregulation of genes involved in cytokine activity and interferon signalling pathway, as well as downregulation of genes involved in oxidative phosphorylation and viral transcription. Host response in COVID-19 subjects for the PBMC group, involved upregulation of genes involved in the complement system and immunoglobulin mediated immune response. *CXCL13, GABRE, IFITM3* were upregulated and *HSPA1B* was downregulated in COVID-19 subjects in both sample groups.

**Conclusion:** Our results indicate the host response to SARS-CoV-2 is compartmentalized and suggests potential biomarkers of response to SARS-CoV-2 infection.

**Highlights:**

- Transcriptomic profiling from publicly available RNA-seq count data revealed a site-specific immune response in COVID-19.
- Host response was found cellular-mediated in nasopharyngeal samples and humoral-mediated in PBMCs samples.
- *CXCL13, GABRE* and *IFITM3* commonly upregulated and *HSPA1B* downregulated in both sample groups highlights the potential of these molecules as markers of response to SARS-CoV-2 infection.

## 1. Introduction

The ongoing global pandemic in 2020, is a viral outbreak of severe acute respiratory syndrome coronavirus 2 (SARS-CoV-2), with over 24 million confirmed cases that account for more than 820,000 confirmed deaths across the world (Pharmaceutical Technology, 2020). SARS-CoV-2 belongs to the coronavirus family (Coronaviridae) of viruses that cause mild or fatal respiratory tract infections in humans. The virus is contagious and infects humans by entering the cells through the attachment of its spike proteins to angiotensin-converting enzyme 2 (ACE2) receptors found on surface of the cell (Mathewson et al., 2008; Shereen et al., 2020; Wang et al., 2020). The clinical manifestations of the disease caused by SARS-CoV-2, coronavirus disease 2019 (COVID-19) include fever, dry cough, breathing difficulties (dyspnoea), headache and fatigue (Lima, 2020; CDC, 2019). Other symptoms observed in some patients are sore throat in the prominent upper respiratory tract, sneezing, anosmia and rhinorrhoea (Huang et al., 2020, Hornuss et al., 2020; Lee et al., 2020; Lechien et al., 2020).

Although there is an expression of interferon (IFN) in other coronavirus infections (Newton *et al*., 2016; Nelemans & Kikkert, 2019), some studies on the immune response to viral infections in addition to the newly understanding of SARS-CoV-2, established that the virus suppresses various mechanisms in the interferon pathway thereby preventing the pattern recognition receptors (PRR), signalling and inhibiting host protein translation (Newton *et al*., 2016; Nelemans & Kikkert, 2019; Lei *et al*., 2020; Mantlo *et al*., 2020). Previous studies also established that triggering of such exuberant inflammatory responses in the host leads to severe lung disease (Channappanavar *et al*., 2016). Most COVID-19 patients with severe clinical presentation and poor prognosis exhibit dysregulated host immune response and enhanced levels of pro-inflammatory cytokines known as the “cytokine storm” (Huang *et al*., 2020). Recent studies done by Haung *et al*. (2020) and Xiong *et al*. (2020) also demonstrated elevated amounts of plasma *IL2, IL6, IL7, IL10, GCSF, IP10, MCP1, MIP1A*, and *TNFα* in severe cases of COVID-19 compared to mild cases, showing inflammatory cytokines release to be significantly associated with COVID-19 progression (Huang *et al*., 2020, Xiong *et al*., 2020).

Transcriptome profiling allows the evaluation of gene expression changes in response to disease or treatment (Voineagu *et al*. 2011). Transcriptome profiling of 8356 cells collected from COVID-19 patients and healthy controls revealed that COVID-19 patients exhibited abnormalities in their bronchoalveolar epithelial cells and that the SARS-CoV-2 virus suppresses gene expression in host cells (He *et al*., 2020). However, the underlying molecular mechanisms of the inflammatory responses and pathogenesis of the COVID-19 in SARS-CoV-2 infection are not clearly understood. Thus, analysing host transcriptional changes in response to SARS-CoV-2 infection is necessary to help describe the biological process underlying the pathogenesis of COVID-19.

In this study, we performed transcriptomic analysis of different samples sources from both COVID-19 patients and healthy controls (Figure 1). We show that transcriptional response to SARS-CoV-2 infection in the two different compartments are distinct and reflect a local inflammation in nasopharyngeal site and circulating immune response in peripheral blood mononuclear cells (PBMC) samples in COVID-19 patients.

**Figure 1.**
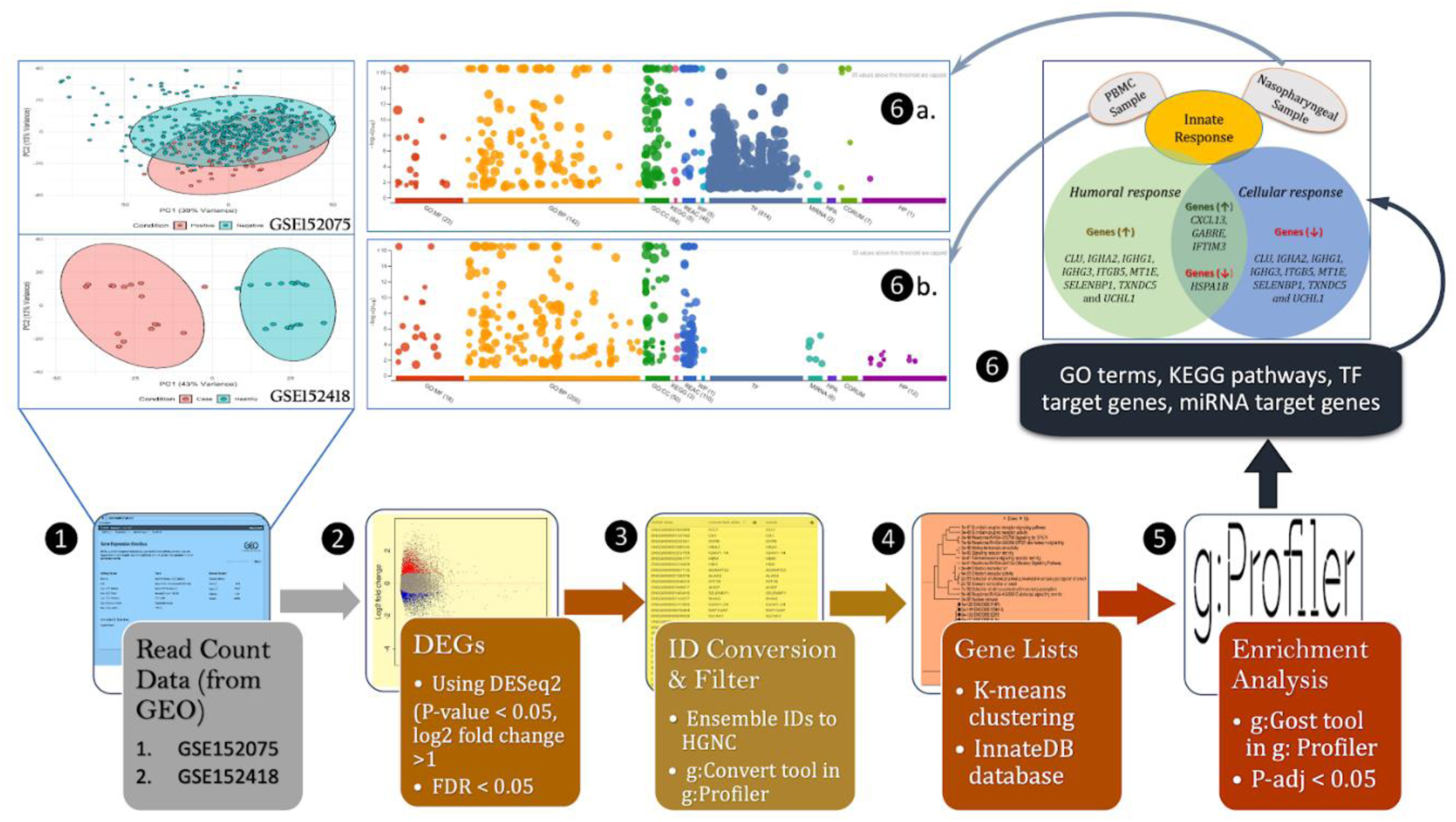
Schematic representation of the workflow in the present study. (1) Gene expression data (RNA-Seq counts) of GSE152418 and GSE152075 datasets were downloaded from the NCBI Gene Expression Omnibus (GEO) database. (2) Differentially expressed gene (DEG) analysis was performed on the control and disease sample groups in each nasopharyngeal and PBMC group using the DESeq2 package in R. Genes that fulfilled the criteria of an adjusted *p*-value < 0.05 and a threshold of log2 fold change > 1 were considered significant. (3) Ensembl IDs of DEGs were converted to HUGO Gene Nomenclature Committee (HGNC) symbols using the g: Convert tool in g: Profiler. (4) An ensemble of hierarchical and k-means clustering was then applied to common DEGs in both the nasopharyngeal and PBMC sample groups and DEGs were also searched for their involvement in innate immunity using the InnateDB database. (5) Functional profiling of DEGs in the two sample groups were carried out using the g:GOSt tool in g: Profiler with adjusted p-value < 0.05 set as cut off for significant terms. (6) Associated gene ontology (GO), molecular function (MF), biological process (BP), KEGG pathway, Reactome pathway, and WikiPathway was identified for each dataset.

## 2. Material and Methods

### 2.1. Study design and datasets

This is a secondary study from publicly available data. Gene expression data (RNA-Seq counts) of GSE152418 and GSE152075 datasets were downloaded from the NCBI Gene Expression Omnibus (GEO) database. The GSE152075 dataset was generated by RNA sequencing of nasopharyngeal swab samples from COVID-19 subjects and healthy controls (Randhawa et al., 2020; Lieberman *et al*., 2020). The dataset was composed of 430 nasopharyngeal swab samples of COVID-19 subjects and 54 of healthy controls. The median age (years) of the COVID-19 patients enrolled in the study was 54 [2 – 98] in which the majority was female (43%; unknown: 12%) (Lieberman *et al*., 2020). The GSE152418 dataset was generated by RNA sequencing of PBMCs from COVID-19 subjects and healthy controls (Arunachalam et al., 2020). The dataset was composed of 16 PBMC samples of COVID-19 subjects and of 17 healthy controls. The median age (years) of the COVID-19 patients was 56 [25 – 94] with a majority of male (55%) (Arunachalam et al., 2020). SARS-CoV-2 infection was confirmed by RT-PCR in both studies. Nasopharyngeal samples were collected in the emergency departments on admission of the patients and PBMC samples during hospitalization.

### 2.2. Differentially expressed gene (DEGs) analysis

DEGs analysis was performed on the control and disease samples in each nasopharyngeal and PBMC group using the DESeq2 package in R (Love et al., 2014). Genes that fulfilled the criteria of an adjusted *p*-value < 0.05 and a threshold of log2 fold change > 1 were considered significant. Ensembl IDs were converted to HUGO Gene Nomenclature Committee (HGNC) symbols using the g: Convert tool in g: Profiler (Raudvere et al., 2019). The common DEGs in both the nasopharyngeal and PBMC samples were identified. An ensemble of hierarchical and k-means clustering was then applied to these common DEGs to determine their discriminatory utility of the samples into their respective sample groups (i.e. Nasopharyngeal or PBMC). The DEGs were also searched for their involvement in innate immunity using the InnateDB database (Breuer et al., 2013).

### 2.3. Gene enrichment analysis

Functional profiling of the upregulated and downregulated DEGs in the two sample groups were carried out using the g:GOSt tool in g: Profiler (Raudvere *et al*., 2019) with adjusted p-value < 0.05 set as cut off for significant terms.

### 2.4. Expression of the results

The results were expressed in tables, graphs and illustrations when convenient. Heatmaps of the top 20 genes (10 up regulated and 10 down regulated) in both datasets were plotted using iDEP.91 server. The hkmeans and dendrogram plot functions were applied to a normalised data matrix of both nasopharyngeal and PBMC samples using software Orange Data Mining (Ljubljana University). Functional analysis was plotted using Prism (GraphPad software, Inc).

## 3. Results

### 3.1. Transcriptome profiling of host response to SARS-CoV-2 infection from nasopharyngeal and PBMC samples

An overview of the gene expression in nasopharyngeal and PBMCs sample groups was observed in the principal component analysis (PCA), which suggests how significant the variation among the gene expression was between the two groups (Figure 2). In nasopharyngeal samples, we observed a separation between healthy controls and COVID-19 subjects along the second principal component, although substantial overlap between groups remained (Figure 2, A). A distinct cluster between healthy controls and cases was observed in the PBMC sample group (Figure 2, B). Next, we performed a differential gene expression analysis relative to matched healthy controls, to explore host response to SARS-CoV-2 infection (Figure 2). In the nasopharyngeal sample group, 745 genes were differentially expressed, of which 166 were upregulated, and 579 were downregulated (Figure 2, C). In the PBMC sample group, 532 genes were differentially expressed, of which 524 were upregulated and 8 were downregulated (Figure 2, D).

**Figure 2:**
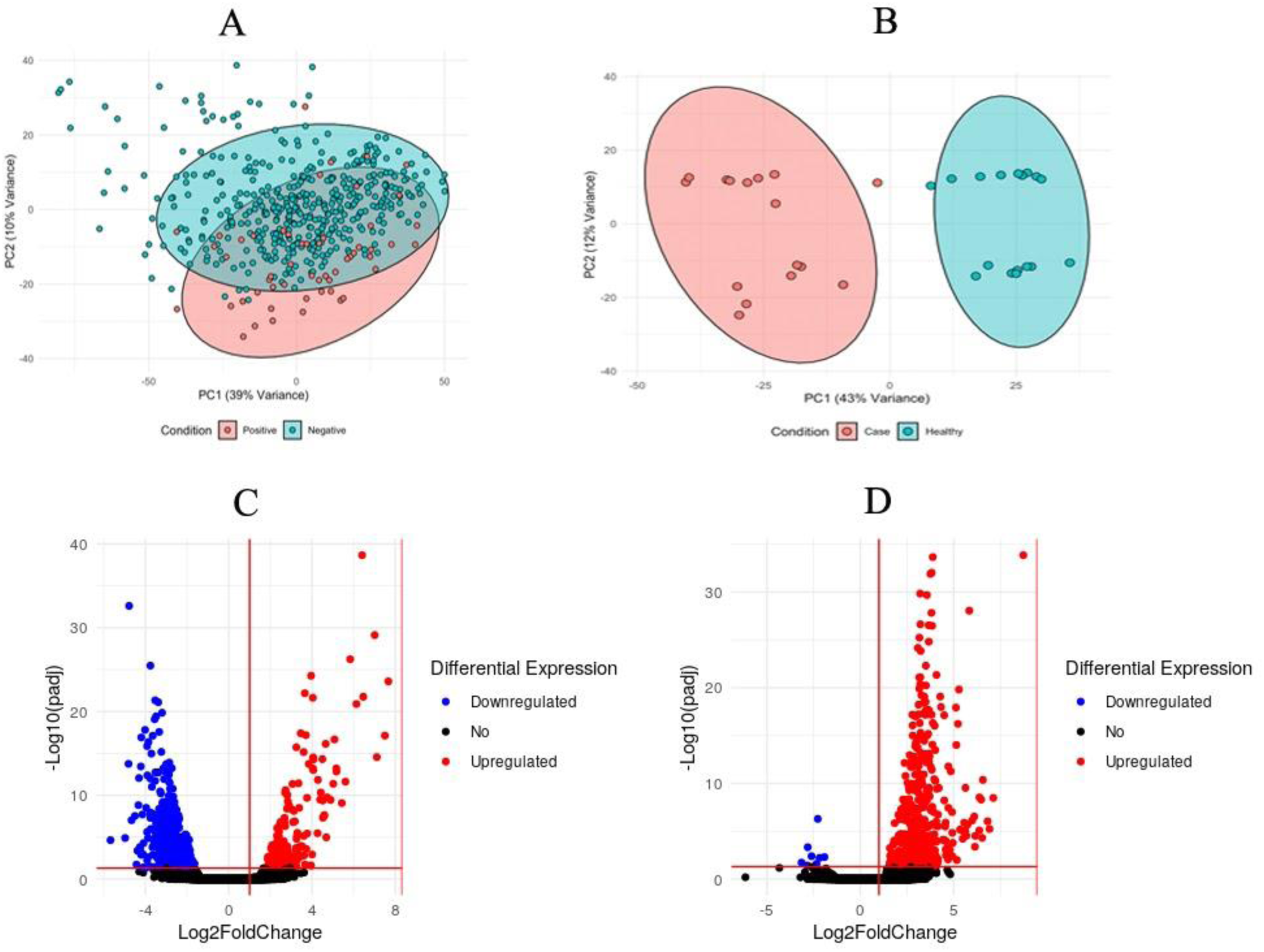
Transcriptomic mapping of nasopharyngeal and PBMC samples in healthy and COVID-19 individuals. Principal component analysis (PCA) from nasopharyngeal samples (A) and PBMC samples (B). Cyan: negative to infection by SARS-CoV-2 (healthy); Pink: positive to infection by SARS-CoV-2 (case). Volcano plots showing the expression of the genes upregulated (red), downregulated (blue) or not differentially expressed (black) in nasopharyngeal samples (C) and PBMCs samples (D).

### 3.2. SARS-CoV-2 infection induces a differential expression pattern in nasopharyngeal and PBMC sample groups

From the number of differentially expressed genes, we next sought to contrast and explore the quantitative changes between the sample groups relative to respective healthy controls. The top significant DEGs evidence a distinct expression pattern in the two sample groups (Figure 3, Tables S1 and S2). Thirteen DEGs including *CLU, CXCL13, GABRE, HSPA1B, IFITM3, IGHA2, IGHG1, IGHG3, ITGB5, MT1E, SELENBP1, TXNDC5* and *UCHL1* were found to be commonly expressed between both sample groups. These genes encode for immunoglobulins, chemokines, ubiquitin, and proteins responsible for stress response and protein degradation. The genes *CXCL13, GABRE* and *IFITM3*, were upregulated in COVID-19 subjects in both sample groups. The genes *CLU, IGHA2, IGHG1, IGHG3, ITGB5, MT1E, SELENBP1, TXNDC5* and *UCHL1* were upregulated in COVID-19 subjects in the PBMC group but downregulated in COVID-19 subjects in the nasopharyngeal sample group. *HSPA1B* was downregulated in both sample groups. The expression pattern of these 13 commonly DEGs consistently discriminated nasopharyngeal and PBMC samples amongst the COVID-19 subjects by hierarchical clustering (Figure 3, C).

**Figure 3.**
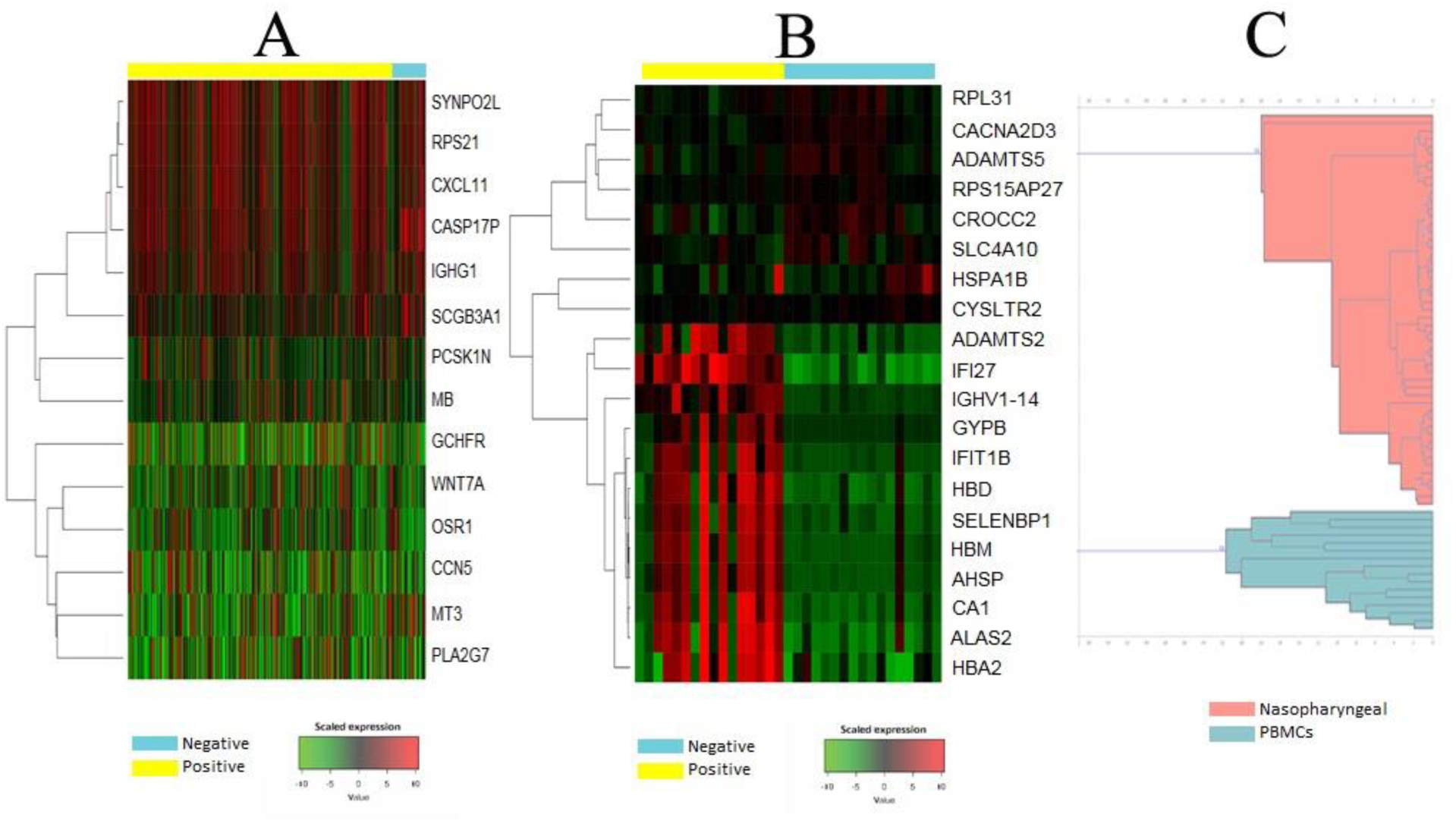
Differential expression of genes in nasopharyngeal and PBMC samples from COVID-19 patients. Heatmaps of gene expression intensity in data series show relative intensity of expression, varying from downregulated (green) to upregulated (red). Shown are the top significant DEGs in healthy donors (negative, blue bar) and COVID-19 patients (positive, yellow bar) in the nasopharyngeal sample group (A) and PBMCs sample group (B). Hierarchical clustering from the common expression of the thirteen genes in nasopharyngeal samples (pink) and PBMCs samples (blue) in patients with COVID-19 (C).

### 3.3. Host immune and metabolic response to SARS-CoV-2 infection

To evaluate biological relevance, we enriched the interaction between the DEGs. Functional profiling of the nasopharyngeal upregulated DEGs identified significant terms such as chemokine activity, cytokine activity, inflammatory response, and type 1 interferon signalling pathway (Figure 4, A), indicating a cellular-mediated immune response. Functional profiling of the upregulated DEGs in COVID-19 subjects in the PBMC group identified significant terms such as immunoglobulin receptor binding, B cell-mediated immunity, complement activation, haemoglobin binding, oxygen carrier activity and cell division (Figure 4, B). The transcriptional profile shown by PBMCs reflects a humoral and complement mediated response against SARS-CoV-2 infection. Ninety-two (92) of the DEGs in the nasopharyngeal group were found to be involved in innate immunity, of which 53 were upregulated, and 39 were downregulated. Functional profiling of the downregulated DEGs in the nasopharyngeal sample identified significant terms such as RNA binding, oxidoreductase activity, cytochrome-c-oxidase activity, oxidative phosphorylation, viral transcription and selenocysteine synthesis (Table S5). There were only 8 downregulated genes in the PBMCs sample group and they were not functionally enriched. Forty-three (43) of the DEGs in the COVID-19 subjects in the PBMC sample group were found to be involved in innate immunity and were all upregulated. Result tables of functional profiling of DEGs are available in the Supplementary Tables S3 to S5. In summary, functional profiling of the DEGs involved in innate immunity in both sample groups identified terms indicating host response to the viral challenge, but distinct pathways of the immune response. For nasopharyngeal sample groups, the events suggest a cellular-mediated immune response, whereas a humoral-mediated response was observed in the PBMC sample group (Figure 5).

**Figure 4.**
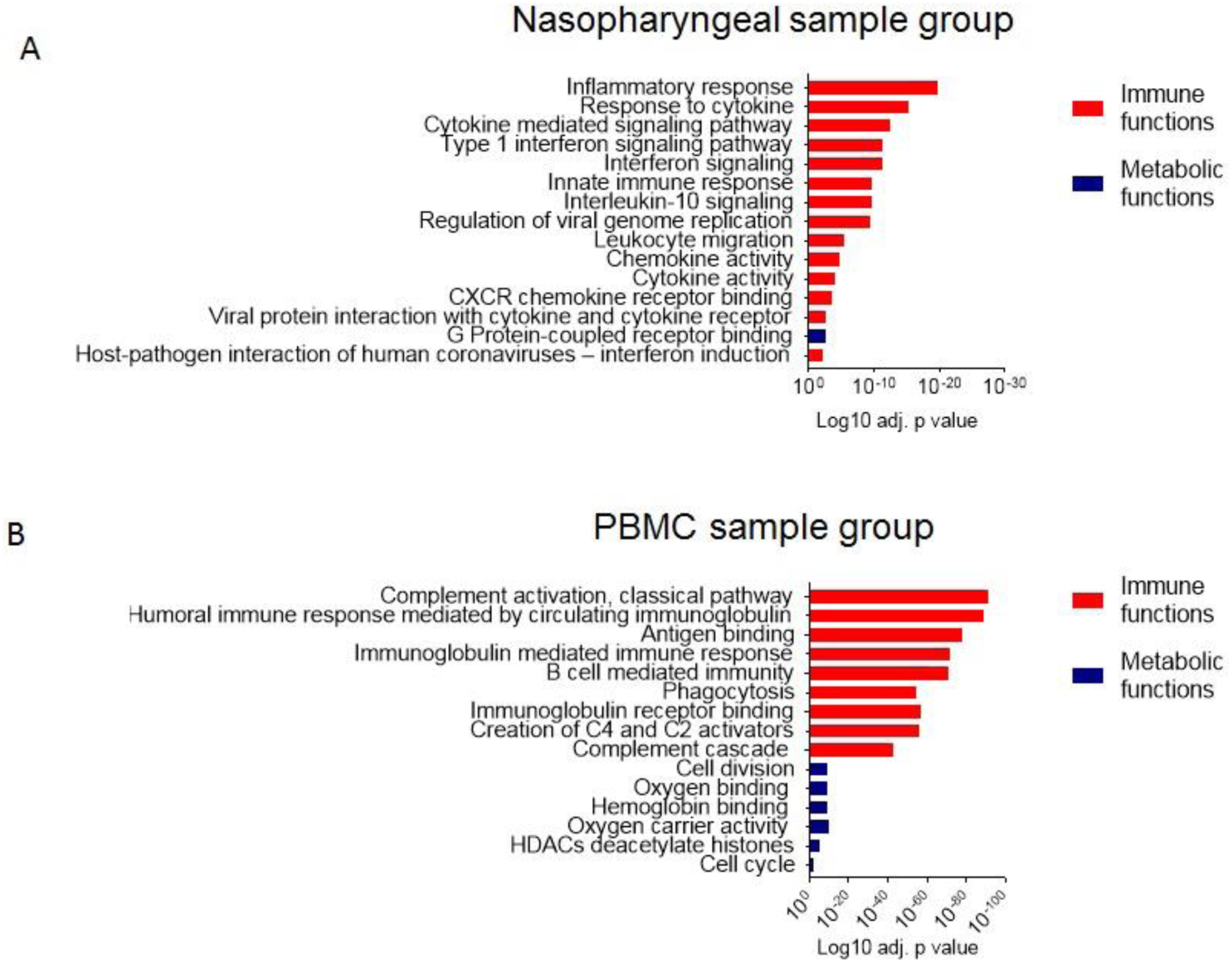
Functional profile of top 15 terms in COVID-19 subjects in nasopharyngeal (A) and PBMC (B) sample groups. Annotations state primarily immune-related functions (red) and metabolic-related functions (blue) played by differentially expressed genes (DEGs) shown in Log10 adjusted p value.

**Figure 5.**
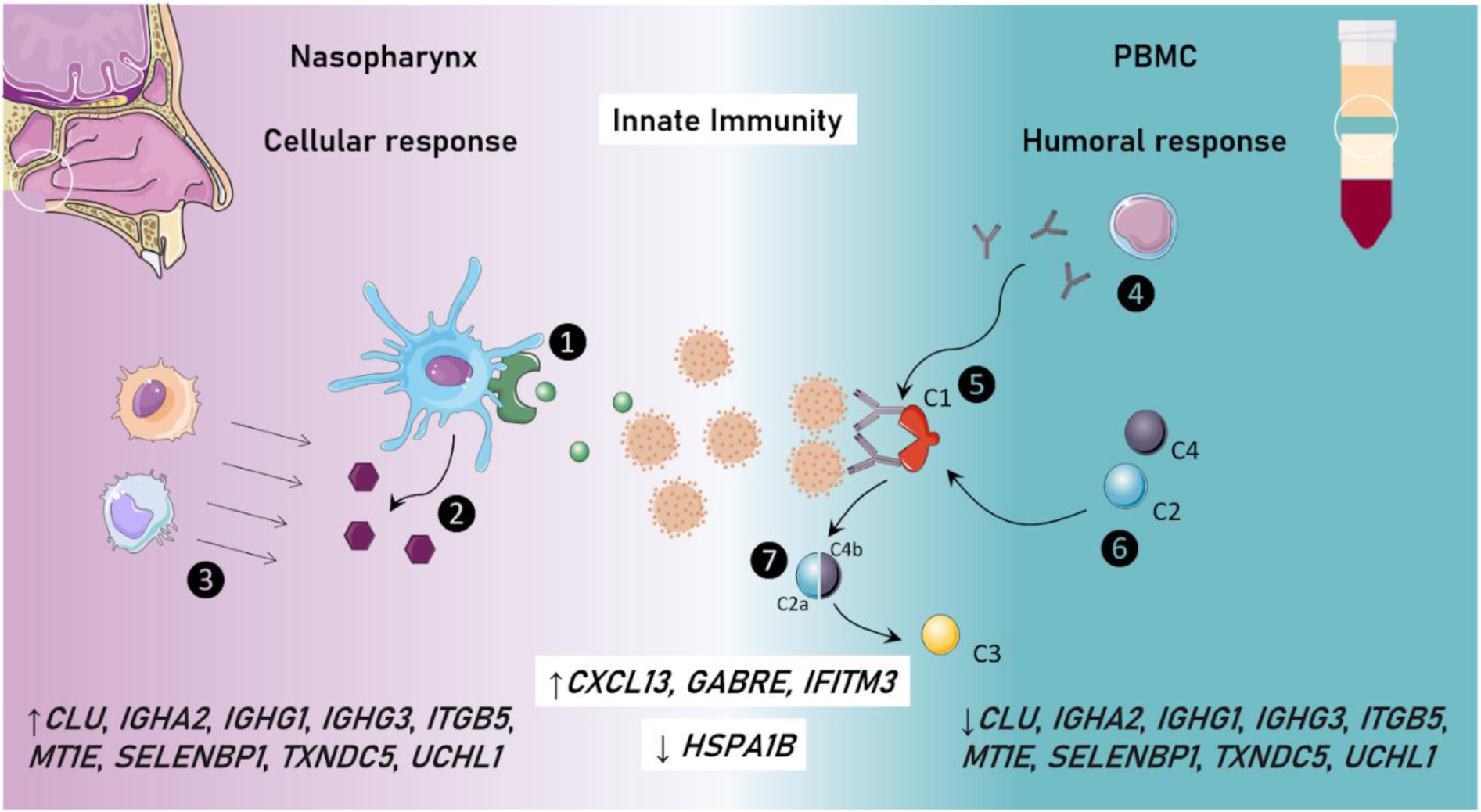
Cellular and molecular alterations in SARS-CoV-2 infection in nasopharyngeal and PBMC samples. Proposed mechanism of the immune response against SARS-CoV-2 infection from functional profiling suggested an innate immunity profile in nasopharyngeal and PBMC samples. Commonly expressed genes appear in upregulation of *CXCL13, GABRE* and *IFITM3* and downregulation of *HSPA1B* for both nasopharyngeal and PBMC samples, downregulation of *CLU, IGHA2, IGHG1, IGHG3, ITGB5, MT1E, SELENBP1, TXNDC5* and *UCHL1* in nasopharyngeal sample and upregulation in PBMC sample. The nasopharyngeal sample revealed an inflammatory response cellular mediated against SARS-CoV-2 infection. Viral protein interaction between cytokine and cytokine receptor occurs (1) in antigen-presenting cells, producing cytokines/chemokines (2) and creating a signalling gradient to favour leukocyte migration (3) to the site of infection. PBMCs sample displayed a humoral response, played by circulating immunoglobulin production mediated by B cells (4) that promote antigen binding, activating the classical pathway of complement system cascade (5). Creation of C4 and C2 activators occurs (6) and cleavages into C2a and C4b (7) to initiate C3 and so on to complete the activation of the complement cascade.

## 4. Discussion

This study was designed to analyse host transcriptional changes in response to COVID-19 from publicly available datasets, to contribute to the knowledge of the biological processes underlying the pathogenesis of COVID-19 and the immune responses elicited. Our results showed a distinct immune response in nasopharyngeal samples and PBMC samples in COVID-19 patients, as evidenced by the differential expression pattern. We have shown that apart from a compartmentalized and site-specific immune response, there was a set of genes commonly regulated in the two sample groups. These observations are in line with previous reports in which SARS-CoV-2 infection induces a highly inflammatory environment observed in nasopharyngeal samples and an innate immune response played by complement system in PBMCs (Blanco-Melo et al., 2020; Xiong et al., 2020).

We showed that nasopharyngeal samples exhibited a regulation for local inflammation, compatible with the tropism of SARS-CoV-2 to nasal epithelial cells (Sungnak et al., 2020). This response was shown to be specific at the infection site, that likely reflect in complementary and systemic elements of immune response in the periphery, as indicated in this study. The infection by SARS-CoV-2 is marked by a systemic inflammatory response that affects various organs in different manners (Gardinassi et al., 2020). Although the elevated inflammatory response may be beneficial in fighting the infection, it may also lead to an over-production of pro-inflammatory cytokines that may cause damage to the olfactory epithelium, as well as adverse outcomes such as alveolar damage, respiratory failure and multiple organ dysfunction (Chen et al., 2017.; Huang et al., 2020; Xiong et al., 2020). Cytokines have been shown to contribute to leukocyte recruitment via activating integrins, promoting migration of adherent leukocytes across endothelium and through the extracellular matrix (Butcher et al., 1996). Therefore, cytokine and chemokine expression in the nasopharynx is possibly associated with the finding of upregulation of leukocytes migration. Blanco-Melo and colleagues (2020), observed a strong chemokine expression in transcriptomic profile from nasopharyngeal swabs and low levels of IFN-I. In contrast to that, through the functional profiling in nasopharyngeal samples, we observed an activity regulated by Type I Interferon. In this study, the soluble levels of IFN-I were not addressed, but a possible counterbalance played by upregulation of anti-inflammatory signalling by IL-10 could be in line to what was observed by Blanco-Melo and colleagues (2020). Hachim *et al*. (2020) also reported that genes, especially the *IFITM3* gene, involved in IFN response to a viral infection such as type 1 interferon signalling pathway, were upregulated in SARS-CoV-2 infected lung epithelial cells. This further elucidates the release of IFN by the innate immune system in response to viral RNAs, such as is the case with SARS-CoV-2 (Nelemans & Kikkert, 2019). Further investigation on the mechanism of recruitment and effector function of responding cells is required in order to better understand the process. The biological processes found in PBMCs are consistently directed towards a humoral-mediated response. Xiong and colleagues (2020) reported similar results from PBMCs, with the observation that genes upregulated in PBMCs of COVID-19 patients were enriched in complement activation, B cell-mediated immunity, and immunoglobulin-mediated humoral immune response. These results indicate a highly specific immune response in periphery. Indeed, Stahel and Barnum (2020) pointed out the importance of the complement system in the pathogenesis of COVID-19. Our results support the need for further studies to uncover the exact mechanism of the complement system in COVID-19.

A molecular signature is observed in the heatmap from PBMCs, albeit not so clearly in nasopharyngeal samples. The sample size, higher in nasopharyngeal sampling, may reflect heterogeneous observations from possible diverse clinical presentations. The fact we have found a specific cellular-mediated immune response in nasopharyngeal samples, even in a heterogeneous group of patients, reinforces the importance of the finding of commonly dysregulated genes with the PBMC sample group. Heat Shock Protein Family A (Hsp70) Member 1B – *HSPA1B* was downregulated in both sample groups. *HSPA1B* codes for a molecular chaperone that plays a vital role in cellular processes including protection of protein from stress and protein quality control system including correct folding of proteins (Kim & Oglesbee, 2012; Scieglinska *et al*., 2019). In addition, it has been found that viral infection stimulates *HSPA1B* the gene to increase its own replication (Kim & Oglesbee, 2012). Therefore, during the viral infection of SARS-CoV-2, downregulation of *HSP1AB* might be an indirect immune response from the host to decrease the viral load by reducing the rate of translation of viral protein molecules. We highlight the need of further research to assess the effect of other RNA viruses on the expression of *HSP1AB* genes and whether the suppression of the gene benefits the host against the viral infection. Chemokine (C-X-C motif) ligand 13 – *CXCL13* preferentially promotes migration of B lymphocytes into follicles. Interferon Induced Transmembrane Protein 3 – *IFITM3* is a protein-coding gene that disrupts the intracellular cholesterol homeostasis and inhibits the entry of viruses to the cell cytoplasm by hindering fusion of the virus with cholesterol depleted endosomes. *IFITM3* is an interferon induced gene that aids in providing immunity against the viral infections (Feeley et al., 2011). Together with downregulated *HSP1AB* and the upregulated genes *CXCL13* and *IFITM3*, we draw attention to these molecules as potential biomarkers of disease progression. *CXCL13* plays a vital role in the migration of B cells during viral infection (Phares *et al*., 2016) and *IFITM3* is also involved in antiviral response (Zhang *et al*., 2020). This may be due to association with the activation of the cytokine storm in infected patients during immune response (Gao *et al*., 2020; Huang *et al*., 2020; Ruan *et al*., 2020). Nevertheless, Hachim *et al*. (2020) also reported an early upregulation of *IFITM3* in SARS-CoV-2 infected patients. Taken together, the roles in the infection played by these commonly expressed genes at a critical site – possibly the entry route of viral-host interaction in the nasopharynx raises the possibility of low invasive tests to investigate these genes in peripheral blood and support personalized medicine.

Here, we provided insights on the potential use of transcriptomic data to address site-specific differential gene expression in response to SARS-CoV-2. Our results direct the potential of markers to discriminate sample source and differential host response to SARS-CoV-2 and possibly other viral infections. Altogether, we suggest further investigation of whether the gene products can be used as potential biomarkers as a COVID-19 screening process.

## 5. Conclusion

Using transcriptomic analysis, we identified differentially expressed genes between COVID-19 subjects in nasopharyngeal and PBMC sample groups in comparison to healthy controls. We also illustrated the prominent immune response genes involved in SARS-CoV-2 infection in both sample types. The differential expression profile observed between the sample groups suggests the existence of different markers of the infection and which may possibly have diagnostic and/or prognostic utility in COVID-19. However, further studies are required to fully characterise the genes involved in the SARS-CoV-2 infection.

## 6. Author contributions

The first three authors have contributed equally. **Caroline Vilas Boas de Melo**^1^: Conceptualization, data curation, formal analysis, investigation, project administration, resources, validation, visualization, writing – original draft, writing – review & editing; **Maruf Ahmed Bhuiyan**^2^: Conceptualization, data curation, formal analysis, investigation, methodology, software, resources, validation, visualization, writing – original draft, writing – review & editing; **Winfred Nyoroka Gatua**^3^: Conceptualization, data curation, formal analysis, investigation, methodology, software, visualization, writing – original draft, writing – review & editing; **Stephen Kanyerezi**^4^: Conceptualization, data curation, methodology, project administration, software, visualization, writing – original draft, writing – review & editing; **Leonard Uzairue**^5^: Conceptualization, project administration, resources, writing – original draft, writing – review & editing; **Priscilla Abechi**^6^: Conceptualization, investigation, project administration, resources, validation, writing – original draft, writing – review & editing; **Karan Kuma**r^7^: Conceptualization, formal analysis, methodology, software, visualization, validation, writing – review & editing; **Jabale Rahmat**^8^: Conceptualization, resources, validation, visualization, writing – original draft, writing – review & editing; **Abdulazeez Giwa**^9^: Conceptualization, data curation, formal analysis, methodology, software, writing – original draft, writing – review & editing; **Gracious Mwandira**^10^: Conceptualization, methodology, visualization, writing – original draft, writing – review & editing; **Abisogun Mujib Olamilekan**^11^: Conceptualization, methodology, visualization, writing – review & editing; **Tiffany Ezinne George**^12^: Conceptualization, investigation, resources, writing – original draft; **Oluwapelumi John Adejinmi**^13^: Conceptualization, writing – original draft, writing – review & editing; **Monsurat Ademidun Ibironke**^14^: Writing – review & editing; **Olayemi David Rotimi**^15^: Conceptualization, resources, writing – original draft; **Dina Aly Mahmoud Aly Abo-Elenein**^16^: Resources, writing – original draft; **Ridwanullah Abiodun Abubakar**^17^: Conceptualization, resources, validation, writing – original draft, writing – review & editing; **Mahmood Usman**^18^: Resources, writing – original draft, writing – review & editing; **Ifeoluwa Adewunmi**^19^: Conceptualization, writing – original draft, writing; **Oyewumi Akinpelu**^20^: Writing – original draft; **Olajide Emmanuel**^21^: Conceptualization, resources; **Khatendra Reang**^22^: Investigation, resources, visualization, writing – original draft; **Akadiri Olalekan**^23^: Conceptualization, resources, writing – original draft, writing – review & editing; **Sarah H. Carl**^24^ : Supervision, writing – review & editing.

## 7. Acknowledgements

We acknowledge HackBio internship, represented by Mr. Adewale Ogunleye, (convenor), Mr. Samson (Head of communication), Mr. Jekayinoluwa (program manager), and Mr. Emmanuel (Head of finance and budget), sponsors and partners of HackBio DNAcompass, CMESBAHF Nigeria, New Life Research Foundation, Galaxy Project, and Biotrust-Scientific. We thank Dr. Washington LC dos-Santos (MD, PhD) for his consultancy in this study. Sarah H Carl, Ph.D., was a mentor for this project.

## Abbreviations

COVID-19: Corona viral disease-2019
SARS-CoV-2: Severe acute respiratory syndrome coronavirus-2
PBMC: Peripheral blood mononuclear cells
PCA: Principal component analysis
GSE: Gene set enrichment
CXCL13: Chemokine ligand 13
GABRE: Gamma-aminobutyric acid receptor subunit epsilon
IFITM3: Interferon (IFN)-induced transmembrane proteins
HSPA1B: Heat shock protein family A (Hsp70) member 1B
IL2: Interleukin-2
IL6: Interleukin-6
IL7: Interleukin-7
IL10: Interleukin-10
GCSF: Granulocyte colony-stimulating factor
IP10: Interferon gamma-induced protein 10
MCP1: Monocyte chemoattractant protein-1
MIP1A: Macrophage inflammatory protein 1-alpha
TNFα: tumor necrosis factor alpha
IFN: Interferon
CLU: Clusterin
IGHA2: Immunoglobulin heavy constant alpha 2
IGHG1: Immunoglobulin heavy constant gamma 1
IGHG3: Immunoglobulin heavy constant gamma 3
ITGB5: Integrin β5
MT1E: Metallothionein 1E
SELENBP1: Selenium binding protein 1
TXNDC5: Thioredoxin domain containing 5
UCHL1: Ubiquitin C-terminal hydrolase L1.

## Supplementary files

**Table S1.**
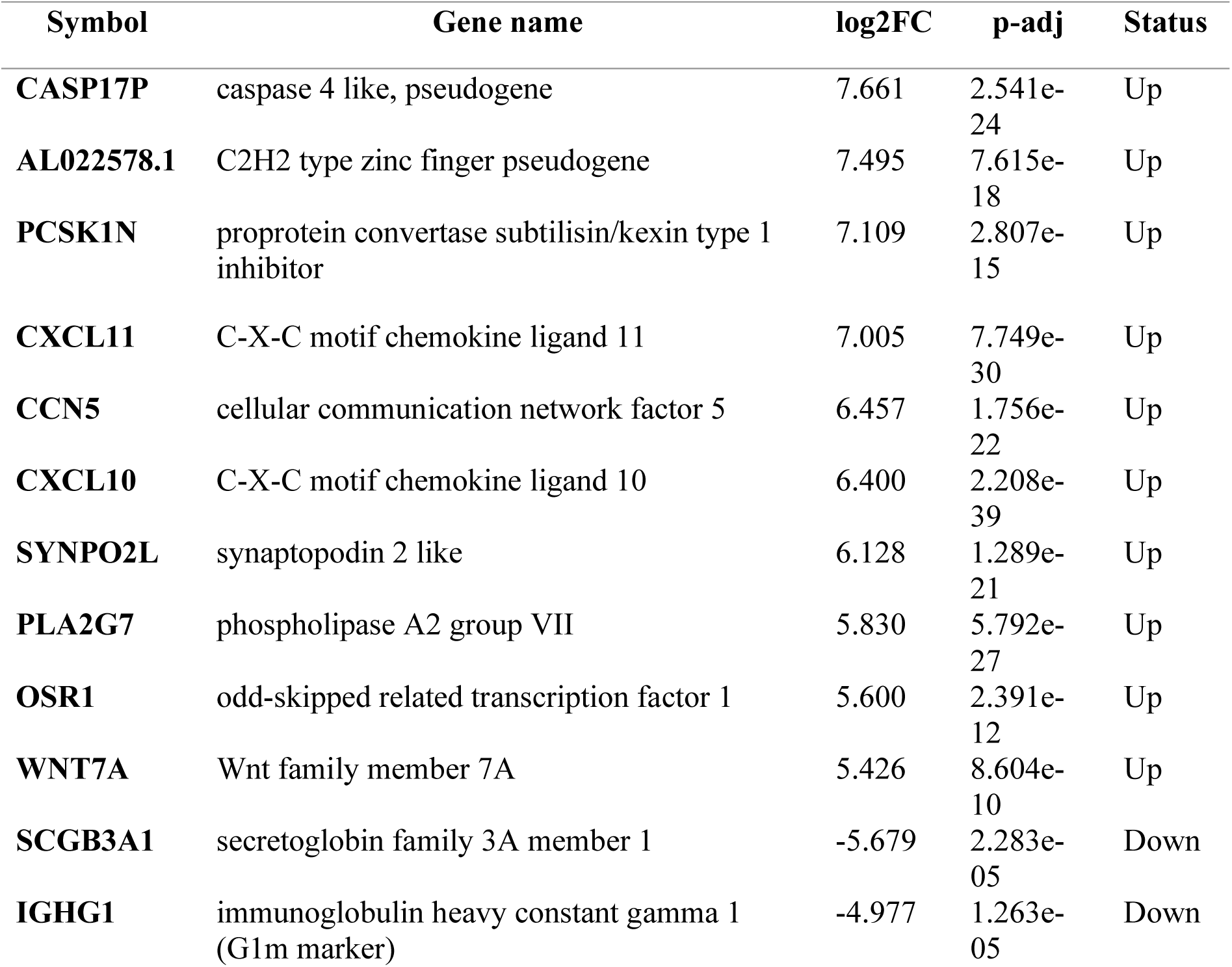
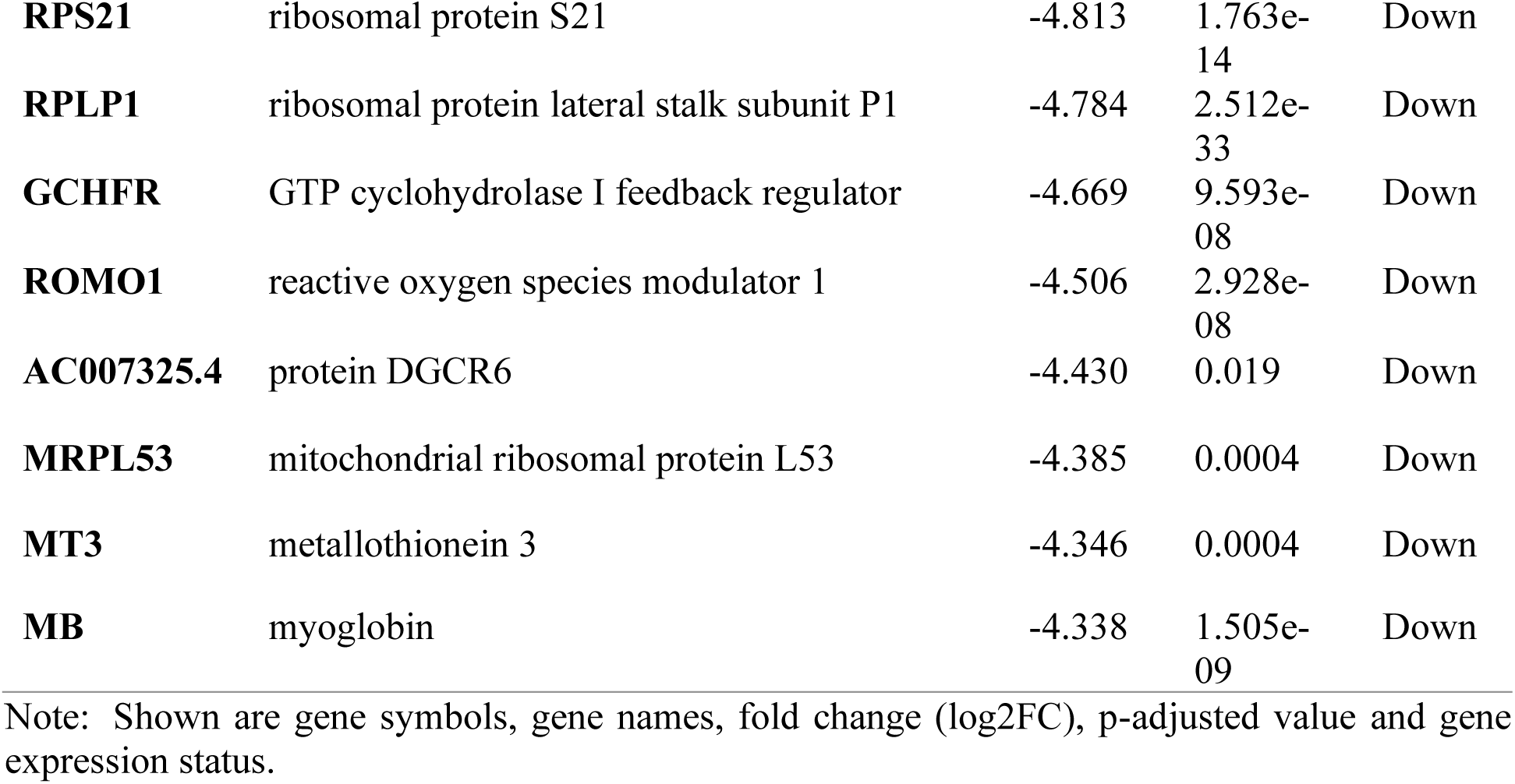
Top significant DEGs (10 Upregulated and 10 Downregulated) in the Nasopharyngeal sample group ordered according to log2FC in disease condition compared to healthy controls.

**Table S2.**
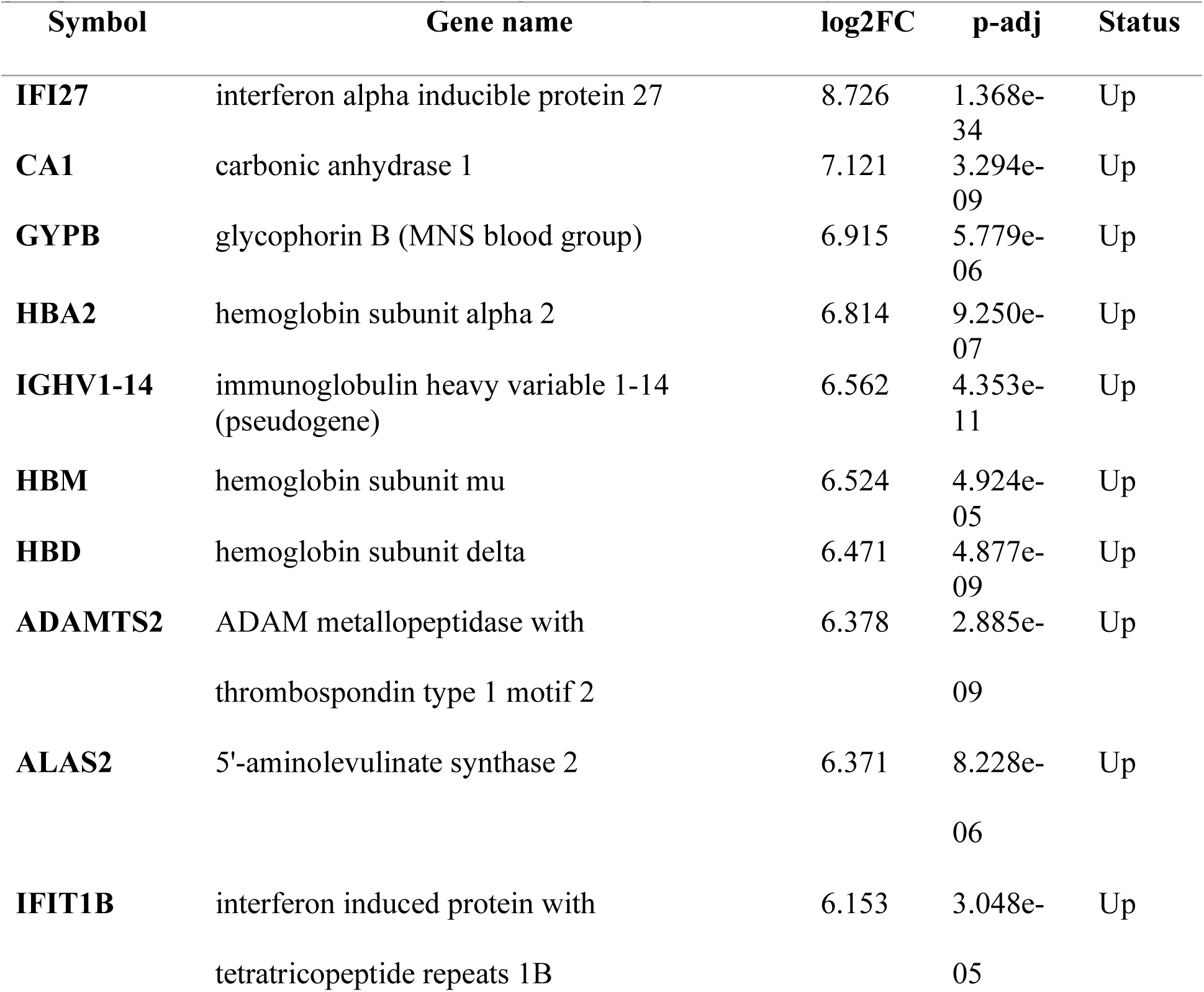
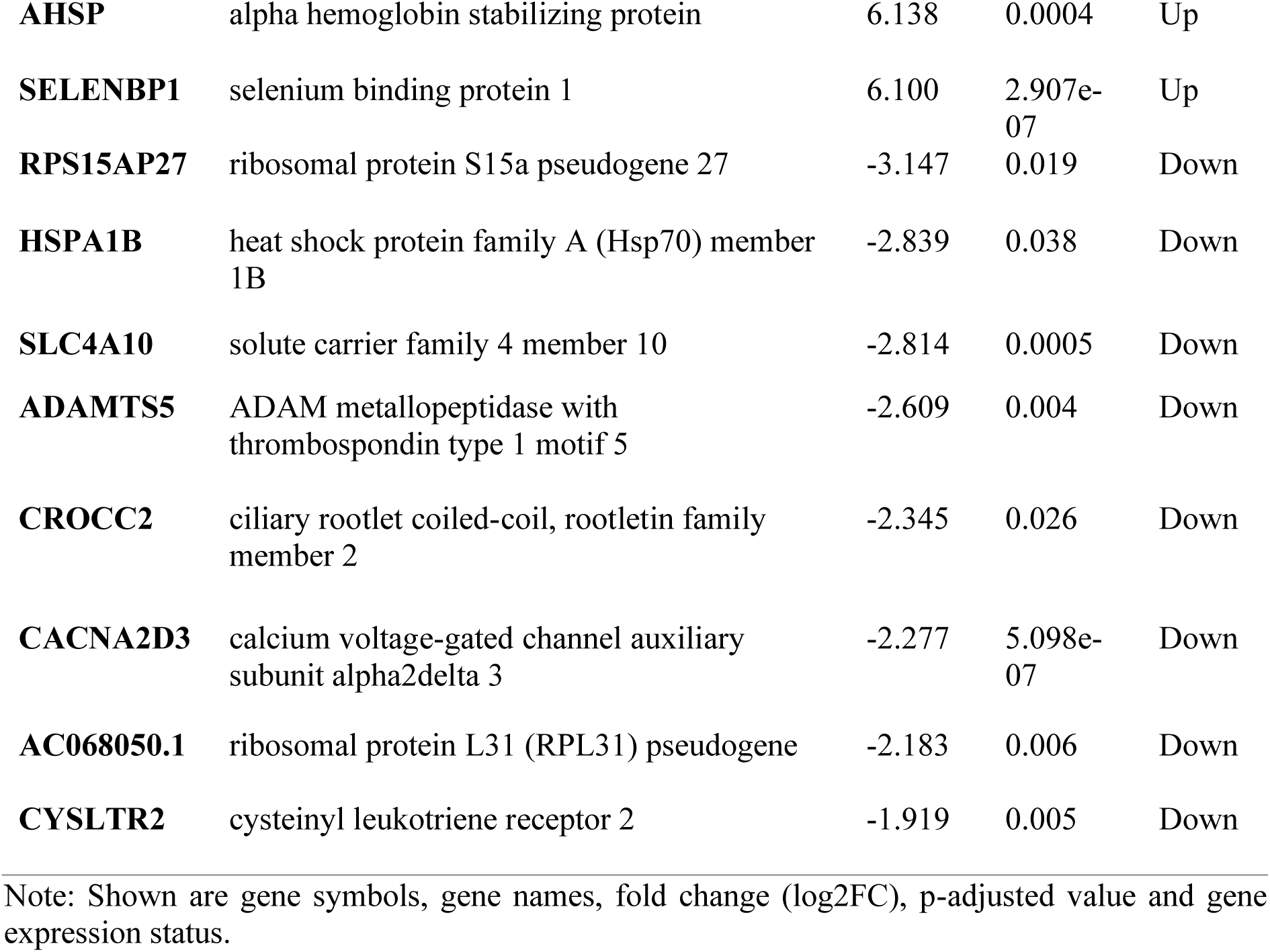
Top significant DEGs (12 Upregulated and 8 Downregulated) in the PBMC sample group in disease condition according to log2FC compared to healthy controls.

**Table S3.**
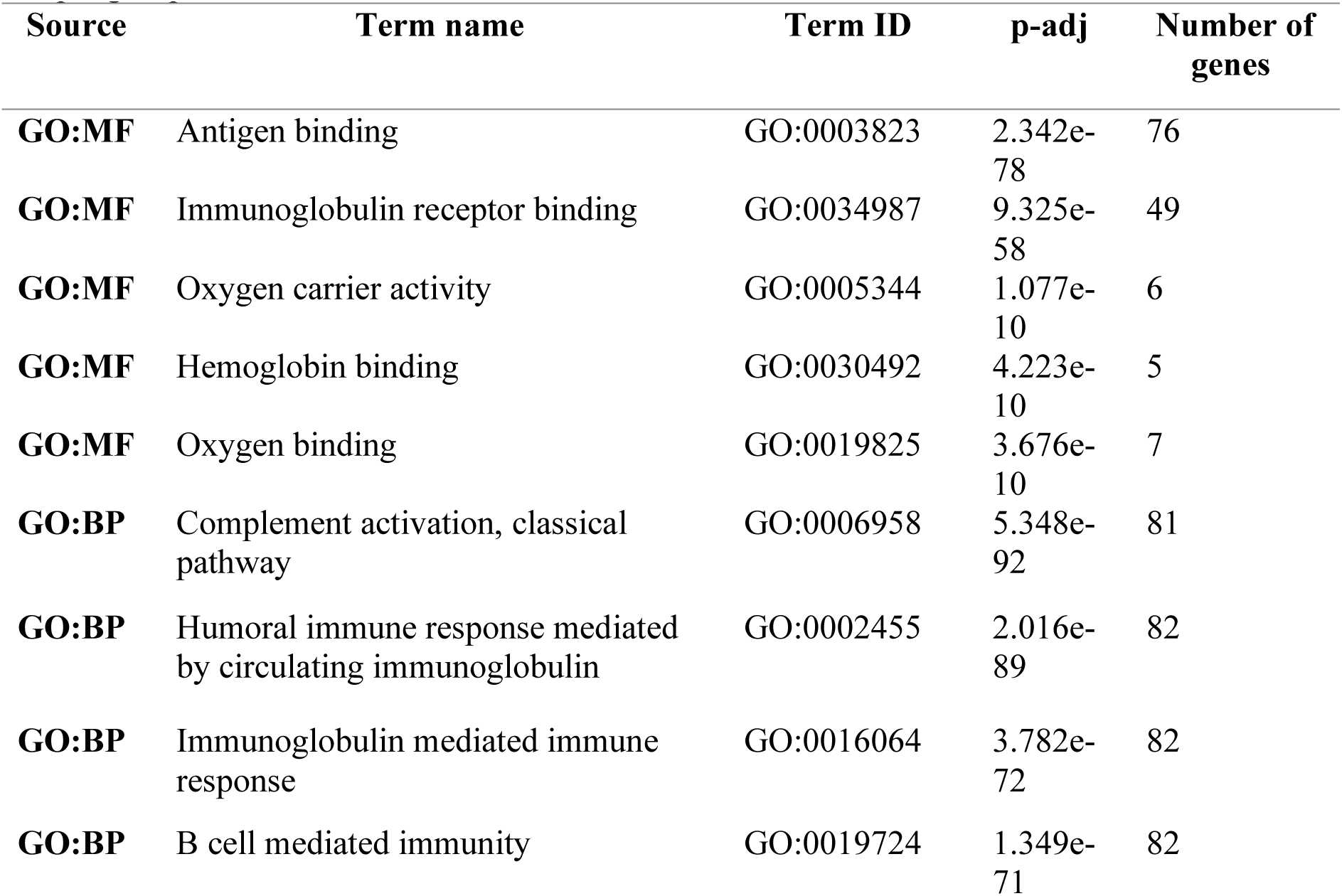
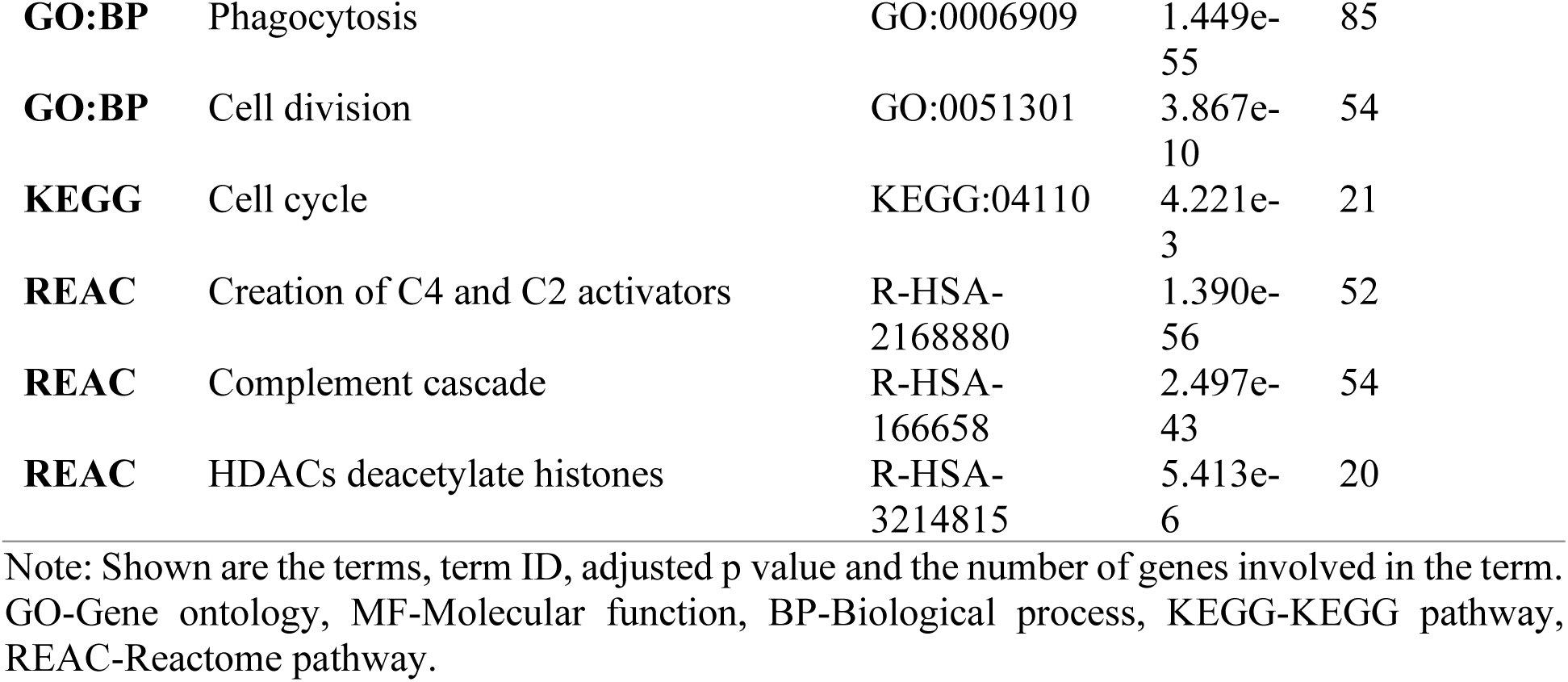
Functional profiling of the upregulated DEGs in the disease condition in the PBMC sample group.

**Table S4.**
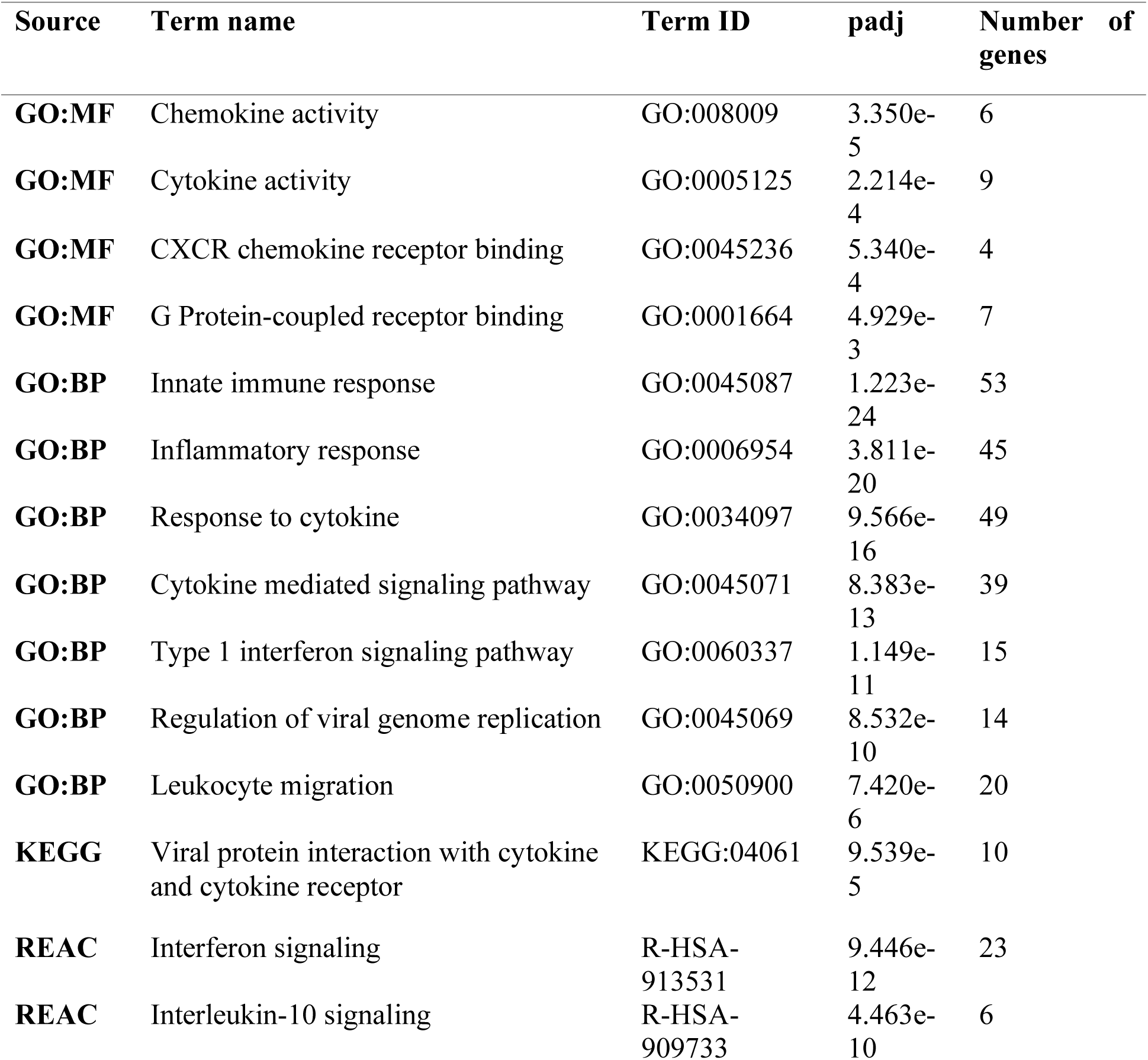
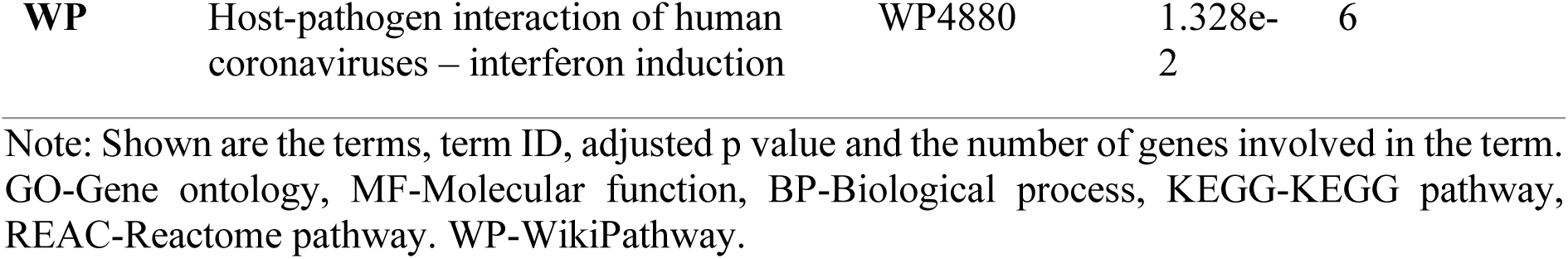
Functional profiling of the upregulated DEGs in the disease condition in the nasopharyngeal sample group.

**Table S5.**
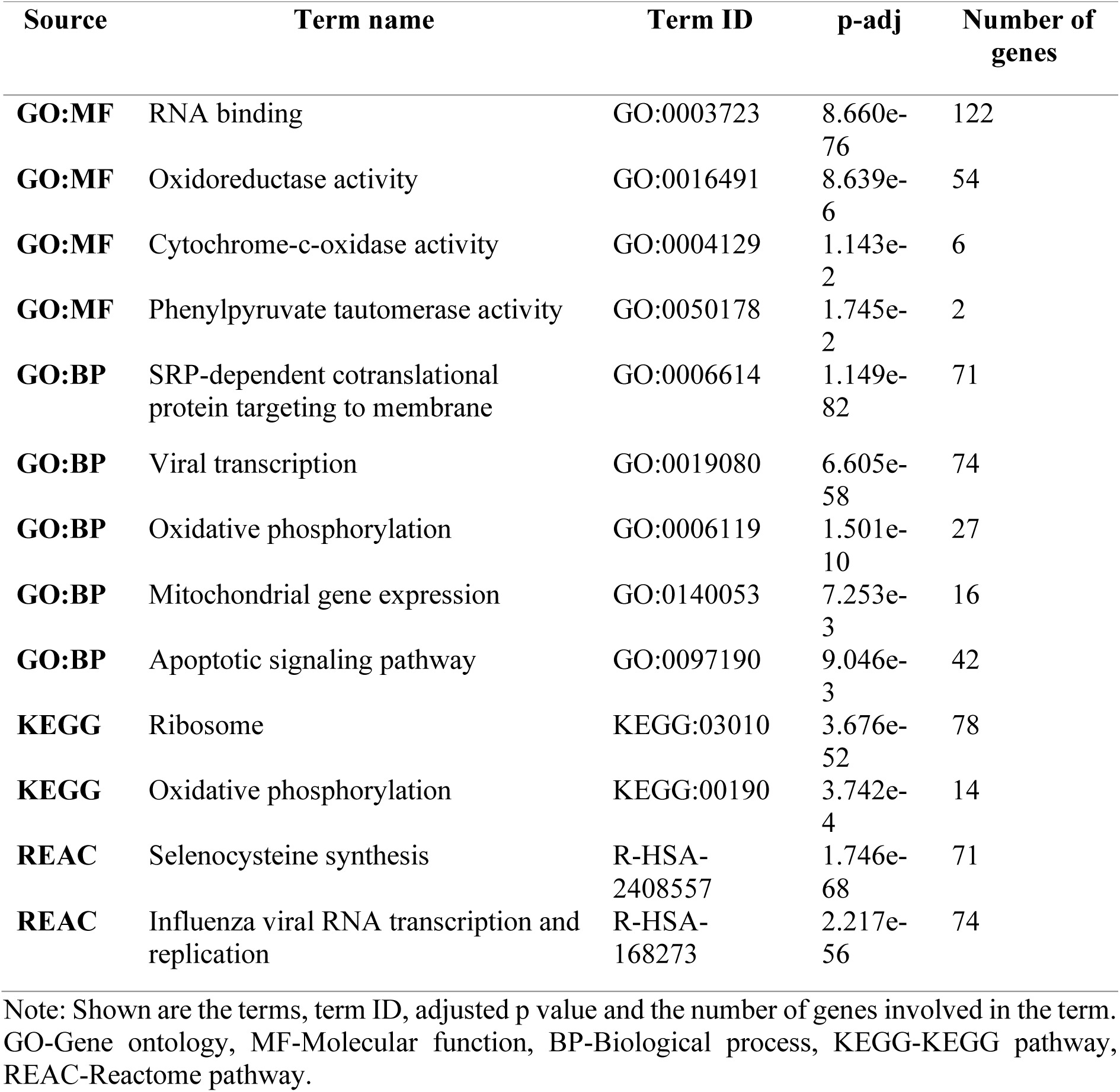
Functional profiling of the downregulated DEGs in the disease condition in the nasopharyngeal sample group.

## References

Arunachalam, P. S., Wimmers, F., Mok, C. K. P., Perera, R. A., Scott, M., Hagan, T., … & Wagh, D. (2020). Systems biological assessment of immunity to mild versus severe COVID-19 infection in humans. Science. https://doi.org/10.1126/science.abc6261

Blanco-Melo, D., Nilsson-Payant, B. E., Liu, W. C., Uhl, S., Hoagland, D., Møller, R., … & Wang, T. T. (2020). Imbalanced host response to SARS-CoV-2 drives development of COVID-19. Cell. https://doi.org/10.1016/j.cell.2020.04.026

Breuer, K., Foroushani, A. K., Laird, M. R., Chen, C., Sribnaia, A., Lo, R., … & Lynn, D. J. (2013). InnateDB: systems biology of innate immunity and beyond—recent updates and continuing curation. Nucleic acids research, 41(D1), D1228–D1233. https://doi.org/10.1093/nar/gks1147

Butcher, E. C., & Picker, L. J. (1996). Lymphocyte homing and homeostasis. Science, 272(5258), 60–67. https://doi.org/10.1126/science.272.5258.60

CDC, (2019). Novel Coronavirus Wuhan, China: Symptoms. CDC. Available at https://www.cdc.gov/coronavirus/2019-ncov/about/symptoms.html

Chen, M., Reed, R. R., & Lane, A. P. (2017). Acute inflammation regulates neuroregeneration through the NF-κB pathway in olfactory epithelium. Proceedings of the National Academy of Sciences, 114(30), 8089–8094. https://doi.org/10.1073/pnas.1620664114

Channappanavar, R., Fehr, A. R., Vijay, R., Mack, M., Zhao, J., Meyerholz, D. K., & Perlman, S. (2016). Dysregulated Type I Interferon and Inflammatory Monocyte-Macrophage Responses Cause Lethal Pneumonia in SARS-CoV-Infected Mice. Cell Host & Microbe, 19(2), 181–193. https://doi.org/10.1016/j.chom.2016.01.007

Feeley, E. M., Sims, J. S., John, S. P., Chin, C. R., Pertel, T., Chen, L. M., Gaiha, G. D., Ryan, B. J., Donis, R. O., Elledge, S. J., & Brass, A. L. (2011). IFITM3 inhibits influenza A virus infection by preventing cytosolic entry. PLoS pathogens, 7(10), e1002337. https://doi.org/10.1371/journal.ppat.1002337

Gao, Y., Li, T., Han, M., Li, X., Wu, D., Xu, Y., Zhu, Y., Liu, Y., Wang, X., & Wang, L. (2020). Diagnostic utility of clinical laboratory data determinations for patients with severe COVID-19. Journal of Medical Virology, 92(7), 791–796. https://doi.org/10.1002/jmv.25770

Gardinassi, L. G., Souza, C. O., Sales-Campos, H., & Fonseca, S. G. (2020). Immune and metabolic signatures of COVID-19 revealed by transcriptomics data reuse. Frontiers in Immunology, 11, 1636. https://doi.org/10.3389/fimmu.2020.01636

Ge, S. X., Son, E. W., & Yao, R. (2018). iDEP: an integrated web application for differential expression and pathway analysis of RNA-Seq data. BMC Bioinformatics, 19(1), 534. https://doi.org/10.1186/s12859-018-2486-6

Hachim, M. Y., Al Heialy, S., Hachim, I. Y., Halwani, R., Senok, A. C., Maghazachi, A. A., & Hamid, Q. (2020). Interferon-Induced Transmembrane Protein (IFITM3) Is Upregulated Explicitly in SARS-CoV-2 Infected Lung Epithelial Cells. Frontiers in Immunology, 11, 1372. https://doi.org/10.3389/fimmu.2020.01372

He, J., Cai, S., Feng, H., Cai, B., Lin, L., Mai, Y., Fan, Y., Zhu, A., Huang, H., Shi, J., Li, D., Wei, Y., Li, Y., Zhao, Y., Pan, Y., Liu, H., Mo, X., He, X., Cao, S., … Chen, J. (2020). Single-cell analysis reveals bronchoalveolar epithelial dysfunction in COVID-19 patients. Protein & Cell, 11(9), 680–687. https://doi.org/10.1007/s13238-020-00752-4

Hornuss, D., Lange, B., Schröter, N., Rieg, S., Kern, W. V., & Wagner, D. (2020). Anosmia in COVID-19 patients. Clinical Microbiology and Infection. https://doi.org/10.1016/j.cmi.2020.05.017

Huang, C., Wang, Y., Li, X., Ren, L., Zhao, J., Hu, Y., Zhang, L., Fan, G., Xu, J., Gu, X., Cheng, Z., Yu, T., Xia, J., Wei, Y., Wu, W., Xie, X., Yin, W., Li, H., Liu, M., … Cao, B. (2020). Clinical features of patients infected with 2019 novel coronavirus in Wuhan, China. The Lancet, 395(10223), 497–506. https://doi.org/10.1016/S0140-6736(20)30183-5

Kim, M. Y., & Oglesbee, M. (2012). Virus-heat shock protein interaction and a novel axis for innate antiviral immunity. Cells, 1(3), 646–666. https://doi.org/10.3390/cells1030646

Lechien, J. R., Chiesa-Estomba MD, C. M., Hans, S., Barillari MD, M. R., Jouffe, L., & Saussez, S. (2020). Loss of Smell and Taste in 2013 European Patients With Mild to Moderate COVID-19. Annals of Internal Medicine. https://doi.org/10.7326/M20-2428

Lee, Y., Min, P., Lee, S., & Kim, S. W. (2020). Prevalence and duration of acute loss of smell or taste in COVID-19 patients. Journal of Korean medical science, 35(18). https://doi.org/10.3346/jkms.2020.35.e174

Lei, X., Dong, X., Ma, R., Wang, W., Xiao, X., Tian, Z., … & Guo, F. (2020). Activation and evasion of type I interferon responses by SARS-CoV-2. Nature Communications, 11(1), 1–12. https://doi.org/10.1038/s41467-020-17665-9

Lieberman, N. A. P., Peddu, V., Xie, H., Shrestha, L., Huang, M.-L., Mears, M. C., Cajimat, M. N., Bente, D. A., Shi, P.-Y., Bovier, F., Roychoudhury, P., Jerome, K. R., Moscona, A., Porotto, M., & Greninger, A. L. (2020). In vivo antiviral host response to SARS-CoV-2 by viral load, sex, and age. BioRxiv, 2020.06.22.165225. https://doi.org/10.1101/2020.06.22.165225

Lima C. (2020). Information about the new coronavirus disease (COVID-19). Radiologia brasileira, 53(2), V–VI. https://doi.org/10.1590/0100-3984.2020.53.2e1

Love, M. I., Huber, W., & Anders, S. (2014). Moderated estimation of fold change and dispersion for RNA-seq data with DESeq2. Genome biology, 15(12), 550. https://doi.org/10.1186/s13059-014-0550-8

Mantlo, E., Bukreyeva, N., Maruyama, J., Paessler, S., & Huang, C. (2020). Antiviral activities of type I interferons to SARS-CoV-2 infection. Antiviral research, 104811. https://doi.org/10.1016/j.antiviral.2020.104811

Mathewson, A. C., Bishop, A., Yao, Y., Kemp, F., Ren, J., Chen, H., Xu, X., Berkhout, B., van der Hoek, L., & Jones, I. M. (2008). Interaction of severe acute respiratory syndrome-coronavirus and NL63 coronavirus spike proteins with angiotensin-converting enzyme-2. Journal of General Virology, 89(11), 2741–2745. https://doi.org/10.1099/vir.0.2008/003962-0

Nelemans, T., & Kikkert, M. (2019). Viral Innate Immune Evasion and the Pathogenesis of Emerging RNA Virus Infections. Viruses, 11(10). https://doi.org/10.3390/v11100961

Newton, A. H., Cardani, A., & Braciale, T. J. (2016). The host immune response in respiratory virus infection: balancing virus clearance and immunopathology. Seminars in Immunopathology, 38(4), 471–482. https://doi.org/10.1007/s00281-016-0558-0

Phares, T. W., DiSano, K. D., Stohlman, S. A., Segal, B. M., & Bergmann, C. C. (2016). CXCL13 promotes isotype-switched B cell accumulation to the central nervous system during viral encephalomyelitis. Brain, Behavior, and Immunity, 54, 128–139. https://doi.org/10.1016/j.bbi.2016.01.016

Pharmaceutical Technology. (2020, August 26). Covid-19 International update: Global Covid-19 cases near 24 million – deaths near 820,000 – more re-infections confirmed. Retrieved from https://www.pharmaceutical-technology.com/special-focus/covid-19/international-update-global-covid-19-cases-near-24-million-deaths-near-820000-more-re-infections-confirmed/

Randhawa, A. K., Fisher, L. H., Greninger, A. L., Li, S. S., Andriesen, J., Corey, L., & Jerome, K. R. (2020). Changes in SARS-CoV-2 Positivity Rate in Outpatients in Seattle and Washington State, March 1–April 16, 2020. Jama, 323(22), 2334–2336. https://doi.org/10.1001/jama.2020.8097

Raudvere, U., Kolberg, L., Kuzmin, I., Arak, T., Adler, P., Peterson, H., & Vilo, J. (2019). g:Profiler: a web server for functional enrichment analysis and conversions of gene lists (2019 update). Nucleic Acids Research, 47(W1), W191–W198. https://doi.org/10.1093/nar/gkz369

Ruan, Q., Yang, K., Wang, W., Jiang, L., & Song, J. (2020). Correction to: Clinical predictors of mortality due to COVID-19 based on an analysis of data of 150 patients from Wuhan, China. Intensive Care Medicine, 46(6), 1294–1297. https://doi.org/10.1007/s00134-020-06028-z

Scieglinska, D., Krawczyk, Z., Sojka, D. R., & Gogler-Pigłowska, A. (2019). Heat shock proteins in the physiology and pathophysiology of epidermal keratinocytes. Cell Stress and Chaperones, 24(6), 1027–1044. https://doi.org/10.1007/s12192-019-01044-5

Shereen, M. A., Khan, S., Kazmi, A., Bashir, N., & Siddique, R. (2020). COVID-19 infection: Origin, transmission, and characteristics of human coronaviruses. Journal of Advanced Research, 24, 91–98. https://doi.org/10.1016/j.jare.2020.03.005

Stahel, P. F., & Barnum, S. R. (2020). Complement Inhibition in Coronavirus Disease (COVID)-19: A Neglected Therapeutic Option. Frontiers in Immunology, 11, 1661. https://doi.org/10.3389/fimmu.2020.01661

Sungnak, W., Huang, N., Bécavin, C., Berg, M., Queen, R., Litvinukova, M., … & Worlock, K. B. (2020). SARS-CoV-2 entry factors are highly expressed in nasal epithelial cells together with innate immune genes. Nature medicine, 26(5), 681–687. https://doi.org/10.1038/s41591-020-0868-6

Voineagu, I., Wang, X., Johnston, P., Lowe, J. K., Tian, Y., Horvath, S., Mill, J., Cantor, R. M., Blencowe, B. J., & Geschwind, D. H. (2011). Transcriptomic analysis of autistic brain reveals convergent molecular pathology. Nature, 474(7351), 380–384. https://doi.org/10.1038/nature10110

Wang, Y., Liu, M., & Gao, J. (2020). Enhanced receptor binding of SARS-CoV-2 through networks of hydrogen-bonding and hydrophobic interactions. Proceedings of the National Academy of Sciences, 117(25), 13967–13974. https://doi.org/10.1073/pnas.2008209117

Xiong, Y., Liu, Y., Cao, L., Wang, D., Guo, M., Jiang, A., Guo, D., Hu, W., Yang, J., Tang, Z., Wu, H., Lin, Y., Zhang, M., Zhang, Q., Shi, M., Liu, Y., Zhou, Y., Lan, K., & Chen, Y. (2020). Transcriptomic characteristics of bronchoalveolar lavage fluid and peripheral blood mononuclear cells in COVID-19 patients. Emerging Microbes & Infections, 9(1), 761–770. https://doi.org/10.1080/22221751.2020.1747363

Zhang, Y., Qin, L., Zhao, Y., Zhang, P., Xu, B., Li, K., Liang, L., Zhang, C., Dai, Y., Feng, Y., Sun, J., Hu, Z., Xiang, H., Knight, J. C., Dong, T., & Jin, R. (2020). Interferon-Induced Transmembrane Protein 3 Genetic Variant rs12252-C Associated With Disease Severity in Coronavirus Disease 2019. The Journal of Infectious Diseases, 222(1), 34–37. https://doi.org/10.1093/infdis/jiaa224

