## Supplementary material for "Transcriptomic dysregulations associated with SARS-CoV-2 infection in human nasopharyngeal and peripheral blood mononuclear cells": Table S1

**Table S1.** Top 20 DEGs (10 Upregulated and 10 Downregulated) in the Nasopharyngeal sample group ordered according to log2FC in disease condition compared to healthy controls.

| Symbol | Gene name | log2FC | p-adj | Status |
| --- | --- | --- | --- | --- |
| CASP17P | caspase 4 like, pseudogene | 7.661 | 2.541e-24 | Up |
| AL022578.1 | C2H2 type zinc finger pseudogene | 7.495 | 7.615e-18 | Up |
| PCSK1N | proprotein convertase subtilisin/kexin type 1 inhibitor | 7.109 | 2.807e-15 | Up |
| CXCL11 | C-X-C motif chemokine ligand 11 | 7.005 | 7.749e-30 | Up |
| CCN5 | cellular communication network factor 5 | 6.457 | 1.756e-22 | Up |
| CXCL10 | C-X-C motif chemokine ligand 10 | 6.400 | 2.208e-39 | Up |
| SYNPO2L | synaptopodin 2 like | 6.128 | 1.289e-21 | Up |
| PLA2G7 | phospholipase A2 group VII | 5.830 | 5.792e-27 | Up |
| OSR1 | odd-skipped related transcription factor 1 | 5.600 | 2.391e-12 | Up |
| WNT7A | Wnt family member 7A | 5.426 | 8.604e-10 | Up |
| SCGB3A1 | secretoglobin family 3A member 1 | -5.679 | 2.283e-05 | Down |
| IGHG1 | immunoglobulin heavy constant gamma 1 (G1m marker) | -4.977 | 1.263e-05 | Down |
| RPS21 | ribosomal protein S21 | -4.813 | 1.763e-14 | Down |
| RPLP1 | ribosomal protein lateral stalk subunit P1 | -4.784 | 2.512e-33 | Down |
| GCHFR | GTP cyclohydrolase I feedback regulator | -4.669 | 9.593e-08 | Down |
| ROMO1 | reactive oxygen species modulator 1 | -4.506 | 2.928e-08 | Down |
| AC007325.4 | protein DGCR6 | -4.430 | 0.019 | Down |
| MRPL53 | mitochondrial ribosomal protein L53 | -4.385 | 0.0004 | Down |
| MT3 | metallothionein 3 | -4.346 | 0.0004 | Down |
| MB | myoglobin | -4.338 | 1.505e-09 | Down |

Note:  Shown are gene symbols, gene names, fold change (log2FC), p-adjusted value and gene expression status.
