## Supplementary material for "Transcriptomic dysregulations associated with SARS-CoV-2 infection in human nasopharyngeal and peripheral blood mononuclear cells": Table S2

**Table S2.** Top 20 DEGs (12 Upregulated and 8 Downregulated) in the PBMC sample group in disease condition according to log2FC compared to healthy controls.

| Symbol | Gene name | log2FC | p-adj | Status |
| --- | --- | --- | --- | --- |
| IFI27 | interferon alpha inducible protein 27 | 8.726 | 1.368e-34 | Up |
| CA1 | carbonic anhydrase 1 | 7.121 | 3.294e-09 | Up |
| GYPB | glycophorin B (MNS blood group) | 6.915 | 5.779e-06 | Up |
| HBA2 | hemoglobin subunit alpha 2 | 6.814 | 9.250e-07 | Up |
| IGHV1-14 | immunoglobulin heavy variable 1-14 (pseudogene) | 6.562 | 4.353e-11 | Up |
| HBM | hemoglobin subunit mu | 6.524 | 4.924e-05 | Up |
| HBD | hemoglobin subunit delta | 6.471 | 4.877e-09 | Up |
| ADAMTS2 | ADAM metallopeptidase with thrombospondin type 1 motif 2 | 6.378 | 2.885e-09 | Up |
| ALAS2 | 5'-aminolevulinate synthase 2 | 6.371 | 8.228e-06 | Up |
| IFIT1B | interferon induced protein with tetratricopeptide repeats 1B | 6.153 | 3.048e-05 | Up |
| AHSP | alpha hemoglobin stabilizing protein | 6.138 | 0.0004 | Up |
| SELENBP1 | selenium binding protein 1 | 6.100 | 2.907e-07 | Up |
| RPS15AP27 | ribosomal protein S15a pseudogene 27 | -3.147 | 0.019 | Down |
| HSPA1B | heat shock protein family A (Hsp70) member 1B | -2.839 | 0.038 | Down |
| SLC4A10 | solute carrier family 4 member 10 | -2.814 | 0.0005 | Down |
| ADAMTS5 | ADAM metallopeptidase with thrombospondin type 1 motif 5 | -2.609 | 0.004 | Down |
| CROCC2 | ciliary rootlet coiled-coil, rootletin family member 2 | -2.345 | 0.026 | Down |
| CACNA2D3 | calcium voltage-gated channel auxiliary subunit alpha2delta 3 | -2.277 | 5.098e-07 | Down |
| AC068050.1 | ribosomal protein L31 (RPL31) pseudogene | -2.183 | 0.006 | Down |
| CYSLTR2 | cysteinyl leukotriene receptor 2 | -1.919 | 0.005 | Down |

Note: Shown are gene symbols, gene names, fold change (log2FC), p-adjusted value and gene expression status.
