## Supplementary material for "Transcriptomic dysregulations associated with SARS-CoV-2 infection in human nasopharyngeal and peripheral blood mononuclear cells": Table S3

**Table S3.** Functional profiling of the upregulated DEGs in the disease condition in the PBMC sample group.

| Source | Term name | Term ID | p-adj | Number of genes |
| --- | --- | --- | --- | --- |
| GO:MF | Antigen binding | GO:0003823 | 2.342e-78 | 76 |
| GO:MF | Immunoglobulin receptor binding | GO:0034987 | 9.325e-58 | 49 |
| GO:MF | Oxygen carrier activity | GO:0005344 | 1.077e-10 | 6 |
| GO:MF | Hemoglobin binding | GO:0030492 | 4.223e-10 | 5 |
| GO:MF | Oxygen binding | GO:0019825 | 3.676e-10 | 7 |
| GO:BP | Complement activation, classical pathway | GO:0006958 | 5.348e-92 | 81 |
| GO:BP | Humoral immune response mediated by circulating immunoglobulin | GO:0002455 | 2.016e-89 | 82 |
| GO:BP | Immunoglobulin mediated immune response | GO:0016064 | 3.782e-72 | 82 |
| GO:BP | B cell mediated immunity | GO:0019724 | 1.349e-71 | 82 |
| GO:BP | Phagocytosis | GO:0006909 | 1.449e-55 | 85 |
| GO:BP | Cell division | GO:0051301 | 3.867e-10 | 54 |
| KEGG | Cell cycle | KEGG:04110 | 4.221e-3 | 21 |
| REAC | Creation of C4 and C2 activators | R-HSA-2168880 | 1.390e-56 | 52 |
| REAC | Complement cascade | R-HSA-166658 | 2.497e-43 | 54 |
| REAC | HDACs deacetylate histones | R-HSA-3214815 | 5.413e-6 | 20 |

Note: Shown are the terms, term ID, adjusted p value and the number of genes involved in the term. GO-Gene ontology, MF-Molecular function, BP-Biological process, KEGG-KEGG pathway, REAC-Reactome pathway.
