## Supplementary material for "Transcriptomic dysregulations associated with SARS-CoV-2 infection in human nasopharyngeal and peripheral blood mononuclear cells": Table S4

**Table S4.** Functional profiling of the upregulated DEGs in the disease condition in the nasopharyngeal sample group.

| Source | Term name | Term ID | p-adj | Number of genes |
| --- | --- | --- | --- | --- |
| GO:MF | Chemokine activity | GO:008009 | 3.350e-5 | 6 |
| GO:MF | Cytokine activity | GO:0005125 | 2.214e-4 | 9 |
| GO:MF | CXCR chemokine receptor binding | GO:0045236 | 5.340e-4 | 4 |
| GO:MF | G Protein-coupled receptor binding | GO:0001664 | 4.929e-3 | 7 |
| GO:BP | Innate immune response | GO:0045087 | 1.223e-24 | 53 |
| GO:BP | Inflammatory response | GO:0006954 | 3.811e-20 | 45 |
| GO:BP | Response to cytokine | GO:0034097 | 9.566e-16 | 49 |
| GO:BP | Cytokine mediated signaling pathway | GO:0045071 | 8.383e-13 | 39 |
| GO:BP | Type 1 interferon signaling pathway | GO:0060337 | 1.149e-11 | 15 |
| GO:BP | Regulation of viral genome replication | GO:0045069 | 8.532e-10 | 14 |
| GO:BP | Leukocyte migration | GO:0050900 | 7.420e-6 | 20 |
| KEGG | Viral protein interaction with cytokine and cytokine receptor | KEGG:04061 | 9.539e-5 | 10 |
| REAC | Interferon signaling | R-HSA-913531 | 9.446e-12 | 23 |
| REAC | Interleukin-10 signaling | R-HSA-909733 | 4.463e-10 | 6 |
| WP | Host-pathogen interaction of human coronaviruses – interferon induction | WP4880 | 1.328e-2 | 6 |

Note: Shown are the terms, term ID, adjusted p value and the number of genes involved in the term. GO-Gene ontology, MF-Molecular function, BP-Biological process, KEGG-KEGG pathway, REAC-Reactome pathway. WP-WikiPathway.
