## Supplementary material for "Transcriptomic dysregulations associated with SARS-CoV-2 infection in human nasopharyngeal and peripheral blood mononuclear cells": Table S5

**Table S5.** Functional profiling of the downregulated DEGs in the disease condition in the nasopharyngeal sample group.

| Source | Term name | Term ID | p-adj | Number of genes |
| --- | --- | --- | --- | --- |
| GO:MF | RNA binding | GO:0003723 | 8.660e-76 | 122 |
| GO:MF | Oxidoreductase activity | GO:0016491 | 8.639e-6 | 54 |
| GO:MF | Cytochrome-c-oxidase activity | GO:0004129 | 1.143e-2 | 6 |
| GO:MF | Phenylpyruvate tautomerase activity | GO:0050178 | 1.745e-2 | 2 |
| GO:BP | SRP-dependent cotranslational protein targeting to membrane | GO:0006614 | 1.149e-82 | 71 |
| GO:BP | Viral transcription | GO:0019080 | 6.605e-58 | 74 |
| GO:BP | Oxidative phosphorylation | GO:0006119 | 1.501e-10 | 27 |
| GO:BP | Mitochondrial gene expression | GO:0140053 | 7.253e-3 | 16 |
| GO:BP | Apoptotic signaling pathway | GO:0097190 | 9.046e-3 | 42 |
| KEGG | Ribosome | KEGG:03010 | 3.676e-52 | 78 |
| KEGG | Oxidative phosphorylation | KEGG:00190 | 3.742e-4 | 14 |
| REAC | Selenocysteine synthesis | R-HSA-2408557 | 1.746e-68 | 71 |
| REAC | Influenza viral RNA transcription and replication | R-HSA-168273 | 2.217e-56 | 74 |

Note: Shown are the terms, term ID, adjusted p value and the number of genes involved in the term. GO-Gene ontology, MF-Molecular function, BP-Biological process, KEGG-KEGG pathway, REAC-Reactome pathway.
